# Diet effects on mouse meiotic recombination: a warning for recombination studies

**DOI:** 10.1101/2021.05.12.443881

**Authors:** Angela Belmonte Tebar, Estefania San Martin Perez, Syong Hyun Nam-Cha, Ana Josefa Soler Valls, Nadia D. Singh, Elena de la Casa-Esperon

## Abstract

Meiotic recombination is a critical process for sexually reproducing organisms. This exchange of genetic information between homologous chromosomes during meiosis is important not only because it generates genetic diversity, but also because it is often required for proper chromosome segregation. Consequently, the frequency and distribution of crossovers are tightly controlled to ensure fertility and offspring viability. However, in many systems it has been shown that environmental factors can alter the frequency of crossover events. Two studies in flies and yeast point to nutritional status affecting the frequency of crossing over. However, this question remains unexplored in mammals. Here we test how crossover frequency varies in response to diet in *Mus musculus* males. We use immunohistochemistry to estimate crossover frequency in multiple genotypes under two diet treatments. Our results indicate that while crossover frequency was unaffected by diet in some strains, other strains were sensitive even to small composition changes between two common laboratory chows. Therefore, recombination is both resistant and sensitive to certain dietary changes in a strain-dependent manner and, hence, this response is genetically determined. Our study is the first to report a nutrition effect on genome-wide levels of recombination. Moreover, our work highlights the importance of controlling diet in recombination studies and may point to diet as a potential source of variability among studies, which is relevant for reproducibility.

## INTRODUCTION

Meiotic recombination, or the exchange of genetic material between homologous chromosomes that occurs during meiosis, has been extensively studied since the early 20th century due to its important role in generating genetic variation and as an essential tool for genetic mapping; later, it was also found that recombination is required for proper chromosome segregation during meiosis in many organisms (Hassold and Hunt 2001; Morgan 1913). Alterations in the number or distribution of crossovers can result in chromosome missegregation and aneuploidy, with implications in fertility and offspring health (Hassold and Hunt 2001; Ottolini et al. 2015). This delicate balance between the selective benefits of genetic variation, proper chromosome segregation and reproductive success has been achieved through a tight regulation of the crossing over process, both at the genetic and the epigenetic levels.

Although the control of crossover number and distribution is a complex and yet not fully understood process, it results in three common observations: 1) each homolog must have at least one crossover (the obligate crossover or crossover assurance) (Dumont 2017; Mather 1937; Pardo-Manuel de Villena and Sapienza 2001); 2) crossovers are not independent, because the occurrence of one interferes with the occurrence of a second one nearby (positive crossover interference) (Muller 1916; Sturtevant 1915); 3) crossover numbers can be maintained in spite of variations in the number of double strand breaks (DSB) they originate from (crossover homeostasis) (Baier et al. 2014; Cole et al. 2012; Hunter 2015). Consequently, the recombination rate is constrained within species, although considerable variation is observed between species (Dumont 2017; Dumont and Payseur 2008; Segura et al. 2013).

Nevertheless, recombination rate still can vary between individuals of the same species: for instance, up to 30% variation has been reported between mice of different strains and hence, of different genetic background (Baier et al. 2014; Dumont 2017; Dumont and Payseur 2011a; Koehler et al. 2002). In addition, crossover frequency is higher in females than in males in both human and mouse, and this difference has been associated to the length of the synaptonemal complex (SC), the proteinaceous scaffold that forms between chromosomes during meiotic prophase and mediates crossover formation (Petkov et al. 2007; Tease and Hulten 2004). A positive correlation between number of crossovers and SC length has also been observed between different inbred strains of mice (Baier et al. 2014; Lynn et al. 2002). These intraspecific studies suggest that genetic differences, as well as chromatin packaging changes, underlie differences in crossover frequency (Baier et al. 2014; de la Casa-Esperon 2012; de la Casa-Esperon and Sapienza 2003; Kleckner 2006; Kleckner et al. 2003). Among the few loci identified in mice that control recombination rate (Balcova et al. 2016; Baudat et al. 2010; de la Casa-Esperon et al. 2002; Myers et al. 2010; Parvanov et al. 2010), *Prdm9* codes for a histone methyltransferase that determines recombination hotspots. These and other studies in several species conclude that recombination is genetically and epigenetically controlled.

In spite of this control, age and certain external factors, such as temperature changes (Bomblies et al. 2015; Lloyd et al. 2018; Plough 1917), diet (Neel 1941), stress (Belyaev and Borodin 1982), toxicants (Gely-Pernot et al. 2017; Susiarjo et al. 2007; Vrooman et al. 2015) and infections (Singh 2019), are capable of modifying the recombination rate. In mice, the best-studied case is that of bisphenol A (BPA) exposure, a component of epoxy resins and polycarbonate plastics used in a wide variety of consumer products. BPA is an endocrine disruptor capable of binding estrogen receptors (Alonso-Magdalena et al. 2012; Susiarjo et al. 2007). Accidental intake of BPA from damaged mouse cages led to the observation in female oocytes of abnormal increases in meiotic disturbances and aneuploidy (Hunt et al. 2003). Subsequent studies showed that BPA exposure also altered the levels of recombination in both female and male meiosis (Brieno-Enriquez et al. 2011; Susiarjo et al. 2007; Vrooman et al. 2015).

From the studies of BPA exposures in mice, we have learnt several lessons: first, the effects on recombination depend on sex and genetic background. BPA exposures resulted in increased recombination in C57BL/6 females, but not in males of the same inbred strain (Susiarjo et al. 2007; Vrooman et al. 2015). However, in males of the CD-1 outbred strain, BPA was able to induce the opposite effect (a reduction of crossover frequency) (Vrooman et al. 2015). Second, BPA also induces other alterations in the germline -e.g., in diverse processes during spermatogenesis, resulting in reduced sperm production (Liu et al. 2013; Wisniewski et al. 2015; Xie et al. 2016). Third, BPA is also capable of eliciting heritable changes and has been associated with epigenetic modifications of the germline (Manikkam et al. 2013; Rahman et al. 2020; Susiarjo et al. 2013; Susiarjo et al. 2015; Wolstenholme et al. 2013; Ziv-Gal et al. 2015).

However, BPA is not unique: other estrogenic substances are also capable of inducing meiosis and recombination changes (Gely-Pernot et al. 2017; Horan et al. 2017; Horan et al. 2018; Vrooman et al. 2015). Interestingly, the meiotic disturbances caused by BPA on metaphase II mouse oocytes can be prevented by a diet rich in phytoestrogens which, in turn, can elicit abnormalities in absence of BPA (Muhlhauser et al. 2009); phytoestrogens are also capable of counteracting methylation changes induced by BPA (Dolinoy et al. 2007). Isoflavone phytoestrogens, mainly genistein and daidzein, are natural compounds abundant in soy and other legumes.

These observations open the question as to whether not just toxicants, but also diets, could affect the recombination rate in mouse. As dietary options could be infinite, we have focused our attention on two categories of diet. First, we were interested in diets that could modify the germline epigenome based on the lessons learnt from BPA studies. Second, we were interested in common diets. With respect to the former, diets that have shown to induce heritable epigenetic changes in the male germline are low-protein, high fat or caloric restriction diets. For instance, in utero 50% caloric restriction can cause metabolic disturbances in the F1 and F2 generations in mice, as well as methylation changes in the transmitting sperm (Martínez et al. 2014; Radford et al. 2014). Male mouse undernourishment can reduce paternal sperm methylation and fertility and have a negative impact on the health of their offspring (Anderson et al. 2006; McPherson et al. 2016). Hence, we decided to test whether paternal undernourishment could affect meiotic recombination rates in a mouse model. This treatment is of particular interest given its relevance to humans, where undernourishment is a burden for many.

With respect to our interest in common diets, it is important to know if these diets have significant effects on recombination and other reproductive phenotypes from a reproducibility perspective. In our animal facility, two diets are regularly used, which differ in their protein, energy and phytoestrogen content (see Materials). As previously discussed, these three dietary factors have been shown to cause meiotic or epigenetic changes in the germline, which leads to the possibility that content differences among common mouse diets may affect recombination as well.

Therefore, we analyzed whether differences between common diets as well as undernourishment can affect recombination rates in adult males. We performed our study in diverse genetic backgrounds, given the variability in crossover frequency, as well as variation in the effects of environmental exposures on recombination and spermatogenesis, reported between different mouse strains (Spearow et al. 1999; Thigpen et al. 2007; Vrooman et al. 2015). We observed that common diets can trigger recombination rate changes in adult male mice. These changes are strain-specific and thus depend on the genetic background. In addition, these diets can elicit sperm motility changes, but no major spermatogenesis disturbances were observed. Therefore, we propose that recombination could be particularly sensitive to certain alterations, potentially epigenetic, caused by diverse effectors such as diet; hence, recombination could be a biomarker of environmentally-induced perturbations in the germline (Donkin and Barres 2018). Moreover, our data compellingly show that diet composition must be taken into account when performing recombination and sperm studies.

## MATERIALS AND METHODS

### Mouse strains and diets

C57BL/6J, PWK/PhJ and MOLF/EiJ mice were obtained from Jackson Laboratory through Charles River and were bred in our facilities for several generations under the same diet and environmental conditions before the studies began. All experimental procedures used in this study were approved by the Committee of Ethics in Animal Care of the University of Castilla-La Mancha. Mouse chow diets were provided by Harlan Laboratories and Capsumlab. Teklad Global 18% Protein Rodent Diet is designed to support gestation, lactation and growth of rodents and, therefore, fed to pregnant and nursing female mice; hence, it will be referred as the “breeding” diet from here on. Teklad Global 14% Protein Rodent Maintenance Diet (and its equivalent Capsumlab Maintenance Complete Chow, used only in the initial set of experiments) is designed to promote longevity and normal body weight in rodents and, therefore, the routinely “maintenance” diet used in many facilities like ours. Description of both diets can be found in Table S1.

We performed two studies: in the initial one, adult males from the three strains were analyzed for the effect of two diets on recombination (undernourishment and breeding diets) provided during 24 days relative to a control group kept *ad libitum* with maintenance diet. Animals switched to breeding diet had free access to the chow, but those of the “undernourishment” group were fed with 50% (2.25 g of maintenance diet) of the regular daily intake (4.5 g, according to Bachmanov et al. 2002). Each diet group had 3 adult mice, except the B6 control and B6 breeding groups, each with 2 mice (average 5.8 months). Health and weight of the animals were regularly monitored. The second study was aimed to verify the differences observed between breeding and maintenance diets in B6 mice and expand the study to testes and sperm phenotypes. Hence, two groups of five B6 mice (average 5.9 months) were fed *ad libitum* during 24 days with each of the two diets. In both studies, all animals were housed in the same room and conditions and treated almost simultaneously, so that each day we processed a mouse of a different treatment group, in order to avoid differences in uncontrolled environmental exposures (temperature, chemicals, etc.) between animals as well as other sources of experimental bias.

Collaborative Cross (CC) founder mice were obtained from the Jackson Laboratory and reared on a different maintenance diet (Laboratory Rodent Diet 5001) in the Biological Research Facility at North Carolina State University. All animals were housed in the same room and were thus subject to the same environmental conditions. At 8 weeks of age, MLH1 immunohistochemistry analysis (see below) was performed in 3 animals per strain (2 in NZO/HILtj) and 25 spermatocytes were analyzed per mouse. All experimental protocols were approved by the Institutional Animal Care and Use Committee of North Carolina State University.

### Tissue collection and processing for histochemistry and sperm analyses

Dates for mouse euthanasia, sample collection and processing were randomized in order to avoid experimental artifacts and the diet group of the processed samples and resulting images were blinded until all measurements were completed to avoid subjective bias during the analyses.

After the 24-day diet period, adult male mice were euthanized by cervical dislocation and weighed. After removing and weighing the testes, chromosome spreads for immunostaining were prepared from one testicle as described below. The other testicle was submerged in Bouin’s solution and processed for histochemistry. Fixed and paraffin-embebed tissues were sectioned and stained with hematoxylin and eosin. Histological analysis of the composition and distribution of the diverse cell types of the seminiferous tubules were performed as previously described (Ahmed and de Rooij 2009; Borg et al. 2010).

Mature spermatozoa were collected from the caudae epididymides in 500 µl modified TYH buffer (in mM: 135 NaCl, 4.7 KCl, 1.7 CaCl_2_, 1.2 KH_2_PO_4_, 1.2 MgSO_4_, 5.6 glucose, 10 HEPES, pH 7.4 adjusted at 37°C with NaOH). Then, the sperm motility was assessed using a computer-aided sperm analyzer (Sperm Class Analyzer^®^ CASA System, Microptic; Barcelona, Spain). Aliquots of 5 µl sperm/sample were placed on a pre-warmed (37°) Leja chamber and examined in a phase contrast microscope (Nikon Eclipse 80i, Tokyo; Japan) equipped with a warmed stage (37°) and a Basler A302fs digital camera (Basler Vision Technologies, Ahrensburg, Germany), which is connected to a computer by an IEEE 1394 interface. Evaluations were made at 10x magnification and at least ten fields or 200 spermatozoa were recorded for each sample. Settings were adjusted to mouse spermatozoa. Recorded parameters were total motility (%), progressive motility (%), curvilinear velocity (VCL, µm/s), straight line velocity (VSL, µm/s), average path velocity (VAP, µm/s), linearity (LIN; %), straightness (STR, %), wobble (WOB; %), lateral head displacement (ALH, µm) and beat cell frequency (BCF, Hz).

Sperm viability was assessed by mixing 5 µl of sperm diluted in THY buffer with 10 µl of eosin-nigrosin for 30 sec and spreading the mix on a slide .The percent of viable sperm was evaluated under the microscope, as eosin stains only the dead sperm, whereas live sperm remains white.

Chromatin stability was assessed using the Sperm Chromatin Structure Assay (SCSA), a flow cytometric test where sperm DNA breaks are evaluated indirectly by analyzing DNA denaturability (Evenson et al. 1980). The assay measures the susceptibility of sperm DNA to acid-induced DNA denaturation, detected by staining with the fluorescent dye acridine orange (AO). Samples were diluted with TNE buffer (0.15 M NaCl, 0.01 M Tris–HCl, 1 mM EDTA; pH 7.4) at a final sperm concentration of 2 x 10^6^ cells and mixed with 400 µl of an acid-detergent solution for 30 seconds. Then, 1.2 ml of AO was added, and samples were evaluated 2 minutes later with a Cytomics FC500 flow cytometer (Beckman Coulter, Brea, CA, USA) AO was excited with a 488 nm argon laser. A total of 5,000 spermatozoa per sample were evaluated. We expressed the extent of DNA denaturation in terms of DNA fragmentation index (DFI), which is the ratio of red to total (red plus green) fluorescence intensity, *i.e.,* the level of denatured DNA over the total DNA. The DFI value was calculated for each sperm cell in a sample, and the resulting DFI frequency profile was obtained. Total DNA fragmentation index (tDFI) was defined as the percentage of spermatozoa with a DFI value over 25. High DNA stainability (HDS), which offers a measure of the percentage of immature sperm cells, was defined as the percentage of spermatozoa with green fluorescence higher than channel 600 (of 1024 channels).

### Immunostaining, microscopy and scoring

Chromosome spreads were prepared from spermatocytes as previously described (Anderson et al. 1999; de Boer et al. 2009; Milano et al. 2019). Briefly, one of the two testes was decapsulated in hypotonic extraction buffer (HEB: 30 mM Tris, pH 8.2, 50 mM sucrose, 17 mM trisodium citrate dihydrate, 5 mM EDTA, 0.5 mM DTT, and 0.5 mM PMSF). Seminiferous tubule fragments were minced in 100mM sucrose and then fixed onto slides with 1% paraformaldehide containing 0.15% Triton X-100 in a humidified chamber. Slides were washed in 1 × PBS with Photo-Flo 200 (Kodak), dried and processed for immunostaining, or stored at -80°C until use.

MLH1 immunostaining allows for identification of about 90% of mammalian crossover sites (Anderson et al. 1999; Cole et al. 2012). For immunostaining, chromosome spreads were washed in 1 x PBS with 0.4% Photo-Flo 200 (Kodak) and 1 x PBS with 0.1% Triton X-100. The slides were blocked in 10% antibody dilution buffer (ADB: 3% bovine serum albumin, 0.05% Triton, 10% goat serum in 1 x PBS). Then, they were incubated overnight at room temperature with primary antibodies: mouse anti-human MLH1 (BD Biosciences) diluted 1:100 and rabbit anti-SCP3 (Abcam) diluted 1:1000 in ADB. Slides were washed as previously and incubated for 1 h at 37° with secondary antibodies: Alexa Fluor 488 goat anti-mouse IgG and goat anti-rabbit conjugated with Alexa Fluor 555 (Life Technologies) diluted 1:1000 and 1:2000 in ADB, respectively. Slides were washed in 0.4% Photo-Flo and mounted with Prolong Gold Antifade Reagent with DAPI (Life Technologies Limited)

All slides were imaged on a Zeiss LSM 710 confocal microscope and analyzed using Zeiss Zen lite software. Only mid-pachytene stage spermatocytes with characteristic sex-body formation (Anderson et al. 1999) and fully synapsed autosomes and XY chromosomes were scored; cells with poor staining or other scoring difficulties were excluded. In the first study, 25 spermatocytes were analyzed per animal, while more (22-40, average 37.3) were examined per mice in the second. For each spermatocyte, we counted the number of foci localizing to the SC of the 19 autosomes. Total SC length was only measured in autosomes, because the appearance and disappearance of foci on the XY bivalent and on the autosomes are temporally uncoupled. Autosomal SC length was initially measured by manually tracing the length of the SYCP3 signal. Given the large dataset of our second experiment, we developed an ImageJ Macro (named “Synaptonemal & CO analyzer”) for SC semiautomatic measuring. This imageJ macro works in four steps: 1) intensity threshold selection of SC signals; 2) automatic detection of each SC, which is reduced to its central skeleton line; 3) manual editing and 4) automatic measuring of the resulting SCs skeletons. For full description, open-access and validation see J. Soriano, A. Belmonte and E. de la Casa-Esperon (in preparation). Distance between MLH1 foci was measured in bivalents with 2 or more foci. The diet group of the samples were blinded until after focus counts and measurements were determined and reviewed by a second observer; any cells with discrepant or ambiguous MLH1 number were discarded.

### Statistical analysis

Comparisons in the average numbers of foci between different strains and/or diets were tested by ANOVA or Student t-test analysis, pooling the results from multiple mice of each group following previous examples (Cole et al. 2012; Vrooman et al. 2015; Zelazowski et al. 2017). Welch ANOVA was applied when homogeneity of variances could not be assumed. These analyses have been successfully employed in comparable studies despite MLH1 foci not following a normal distribution, because of the robustness of ANOVA analysis (Baier et al. 2014; Dumont 2017). Similar conclusions about statistical significance were obtained if non-parametric tests were performed. For statistically significant differences (*P* < 0.05), a Tukey’s post-hoc honestly significant difference (HSD) test was performed to infer which groups differed. A Chi-square test was used to determine significance in the number of bivalents classified according to their foci number (E0-E3) between diet groups. Weight, sperm count and SCSA data were analyzed by Student t-test. Total motility, progressive motile spermatozoa, VCL, VSL, VAP, LIN, ALH, BCF and sperm viability were evaluated by a factorial ANOVA in mice fed with different diets. When the variables were significant (P < 0.05), *post hoc* comparisons with Bonferroni correction were carried out. Analyses were performed using SPSS Statistics software.

We also used a generalized linear model to compare the average numbers of foci between different strains and/or diets. The full model includes effects of strain, diet, animal, and interaction effects. This was implemented in JMP Pro version 14.

## RESULTS

### Recombination levels depend on the genetic background

Variation in crossover frequency, as well as variability in the effects of chemical exposures on recombination, have been observed among mouse strains (Baier et al. 2014; Dumont and Payseur 2011b; Koehler et al. 2002; Vrooman et al. 2015). In order to investigate if diets have an impact on recombination levels, we selected three mouse inbred strains of diverse genetic background: C57BL/6J (B6), PWK/PhJ (PWK) and MOLF/EiJ (MOLF). B6 is a classical inbred strain widely used in recombination studies (Baier et al. 2014; Dumont and Payseur 2011b; Koehler et al. 2002), which is mostly of *Mus musculus domesticus* origin (93% of autosomal sequences (Yang et al. 2011)); PWK was derived from wild mice of *M. m. musculus* subspecies (Gregorova and Forejt 2000) (94% of autosomal sequences of *M. m. musculus* origin according to Yang et al. (2011); MOLF is representative of the Japanese *M. m. molossinus* subspecies, which is the result of the hybridization between *M. m. musculus* and *M. m. castaneus* (Silver 1995).

Our first goal was to determine the baseline crossover frequencies of the three strains with regular maintenance diet. Our analysis reveals a significant effect of strain on MLH1 focus count (*P* << 0.0001, ANOVA). We observed that while C57BL/6 males have 23.54±1.97 MLH1 foci per spermatocyte (mean±SD), MOLF males have significantly more foci per spermatocyte (24.85±1.97; *P* = 0.015, HSD). MLH1 focus counts are substantially higher in PWK males than both C57BL/6 and MOLF males (29.15±2.87; *P* << 0.0001, both comparisons HSD) (Figures 1A and S1 and Table 1). Hence, genetic differences between the selected strains have an impact on the levels of recombination.

**Figure 1.**
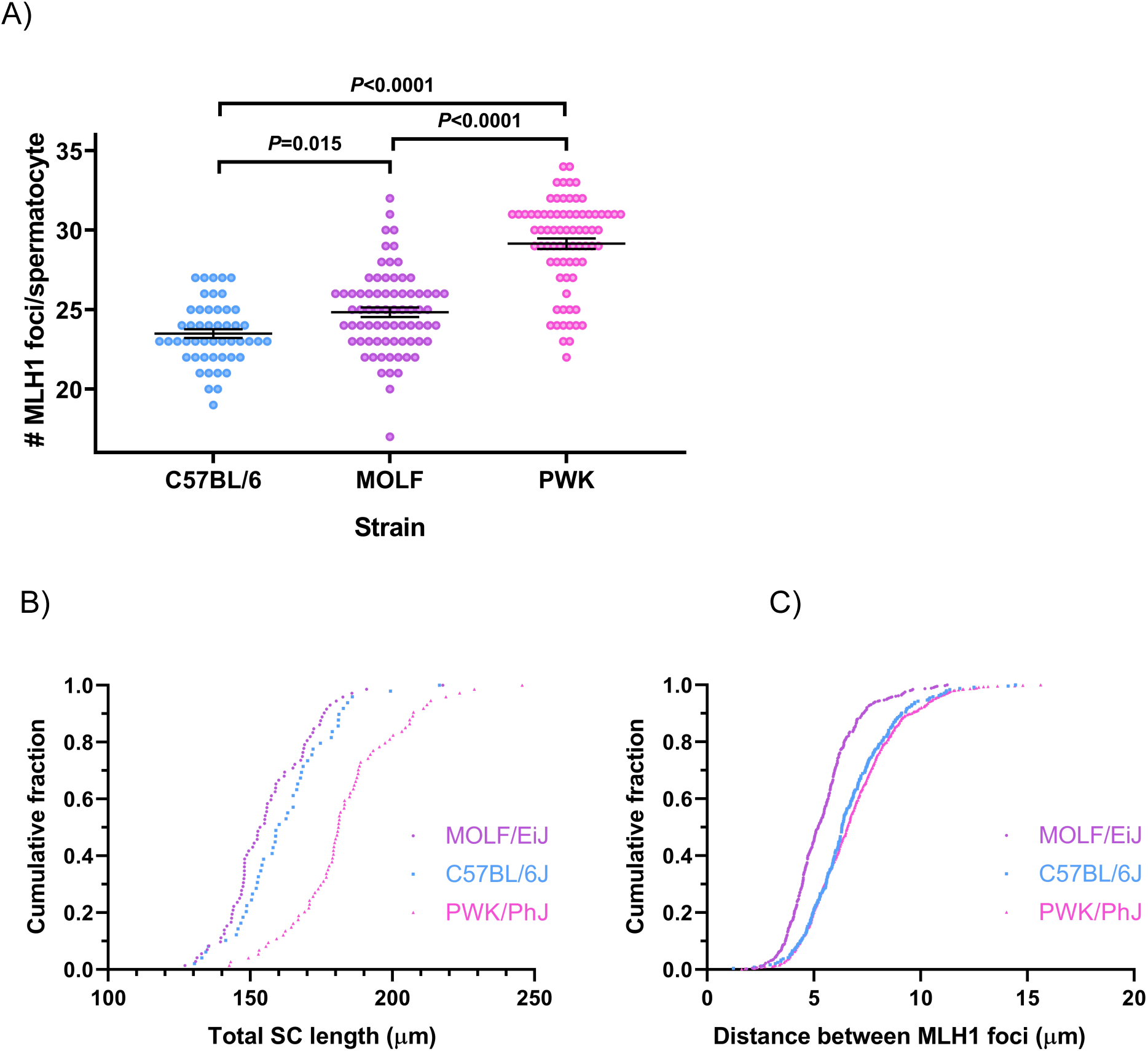
Strain effects on recombination rates, SC length and interference. Recombination levels depend on the genetic background. Autosomal MLH1 foci counts in mid-pachytene spermatocytes are shown in A for mice of the 3 strains fed *ad libitum* with maintenance diet. Significant differences were observed between the three strains, as explained in Table 1. Each dot represents the focus count of a single nucleus. Black bars represent means ± SEM. The cumulative fraction of the total autosomal SC length and the intercrossover distances measured in micrometers are represented in B and C, respectively. PWK spermatocytes have significantly longer SC, while MLH1 interfocus distances in MOLF are significantly shorter than those of the other two strains (Table 1).

**Table 1.** Strain effects on the number of autosomal MLH1 foci and total autosomal SC length per pachytene spermatocyte, and on the length of the SC between MLH1 foci. Comparison of the three strains fed with control maintenance diet shows significant differences in the average number of MLH1 foci (F = 81.5, *P* << 0.0001), SC length (F = 45.4, *P* << 0.0001) and intercrossover distance (F = 73.4, *P* << 0.0001). Post-hoc analysis reveals significant differences in MLH1 foci frequency between the three strains (ab, *P*=0.015; bc, *P* << 0.0001; ac, *P* << 0.0001). In addition to displaying the highest crossover frequency, PWK has the longest SC length (de, *P* << 0.0001 when compared to any of the two other strains). In contrast, MOLF has the shortest interfocus distance *per se* (fg, *P* << 0.0001 when compared to any of the two other strains) or calculated as a percentage of SC length of the corresponding bivalent (hi, *P* << 0.0001 again when MOLF was compared to any of the two other strains). Data were obtained from 50 B6, 75 MOLF and 75 PWK spermatocytes, with 244, 455 and 765 interfocus distance measurements respectively. Analyses were performed by ANOVA and significant differences between groups were assessed using Tukey’s post-hoc tests. Values are shown as mean ± SD.

| Strain | B6 | MOLF | PWK |
| --- | --- | --- | --- |
| MLH1 foci: | 23.54±1.97<sup>a</sup> | 24.85±2.57<sup>b</sup> | 29.15±2.87<sup>c</sup> |
| SC length (µm): | 162.2±17.1<sup>d</sup> | 156.6±16.0<sup>d</sup> | 183.3±19.4<sup>e</sup> |
| Interfocus distance (µm): | 6.63±2.00<sup>f</sup> | 5.43±1.57<sup>g</sup> | 6.83±2.06<sup>f</sup> |
| %Interfocus/SC length: | 64.0±12.9<sup>h</sup> | 56.2±12.8<sup>i</sup> | 63.1±13.3<sup>h</sup> |
| µmSC/MLH1 foci | 6.9 | 6.3 | 6.3 |

### Recombination variability in genetically diverse mice: the Collaborative Cross and Diversity Outbred stock founder strains

C57BL/6 is one of the most widely used mouse strains, but interest for PWK mice, at the other extreme of the crossover frequency, is increasing as a representative strain of the *M. m. musculus* subspecies in many genetic studies and resources, such as the Collaborative Cross (CC) and Diversity Outbred (DO) population. Both resources are the result of crosses between eight founder strains that include B6 and PWK, but also A/J, 129S1/SvImJ, NOD/LtJ, NZO/HlLtJ, CAST/EiJ and WSB/EiJ. These eight strains were selected because they capture most of the genetic diversity present in *Mus musculus* and, therefore, CC and DO mice have become instrumental for multiple genetic studies (Chesler et al. 2008; Churchill et al. 2004; Collaborative Cross Consortium 2012; Roberts et al. 2007; Svenson et al. 2012; Threadgill et al. 2011). Indeed, analysis of CC mice has enabled the characterization of several loci and mechanisms that control recombination (Liu et al. 2014). However, the crossover rate had not been described for all the founder strains (including PWK). Therefore, we decided to characterize the crossover frequency of the eight strains under the same developmental and environmental conditions for the benefit of two purposes: first, for future genetic studies with the CC and DO mice, including further analyses of loci involved in recombination; and second, for comparing the crossover variation of the strains selected for our study relative to the extent of recombination variability present in *Mus musculus*.

We observe 22.21±1.86 MLH1 foci per spermatocyte in CAST/EiJ mice, 22.80±2.22 in NZO/HlLtJ, 22.89±2.60 in C57BL/6J, 23.40±2.35 in A/J, 23.75±2.61 in 129S1/SvImJ, 24.19±23.36 in WSB/EiJ, 25.48±2.40 in NOD/LtJ, 28.11±3.83 in PWK/PhJ (Figure 2). There are significant differences in the MLH1 foci per spermatocyte between the strains (*P* << 0.0001, ANOVA) although, as shown in Figure 2, the majority has frequencies similar to that of B6. CAST/EiJ has the least MLH1 foci count per spermatocyte, but not significantly lower than B6. At the other extreme, PWK values are again significantly higher than any of the other strains (*P* < 0.0001 in all cases), with 27% more crossovers than the CAST/EiJ spermatocytes, a variability in crossover frequency between mice of different strains similar to that reported in other studies (Baier et al. 2014; Dumont and Payseur 2011b; Koehler et al. 2002). Intermediate values are observed in NOD/LtJ (also significantly higher than B6 (*P* < 0.0001)) and WSB/EiJ.

**Figure 2.**
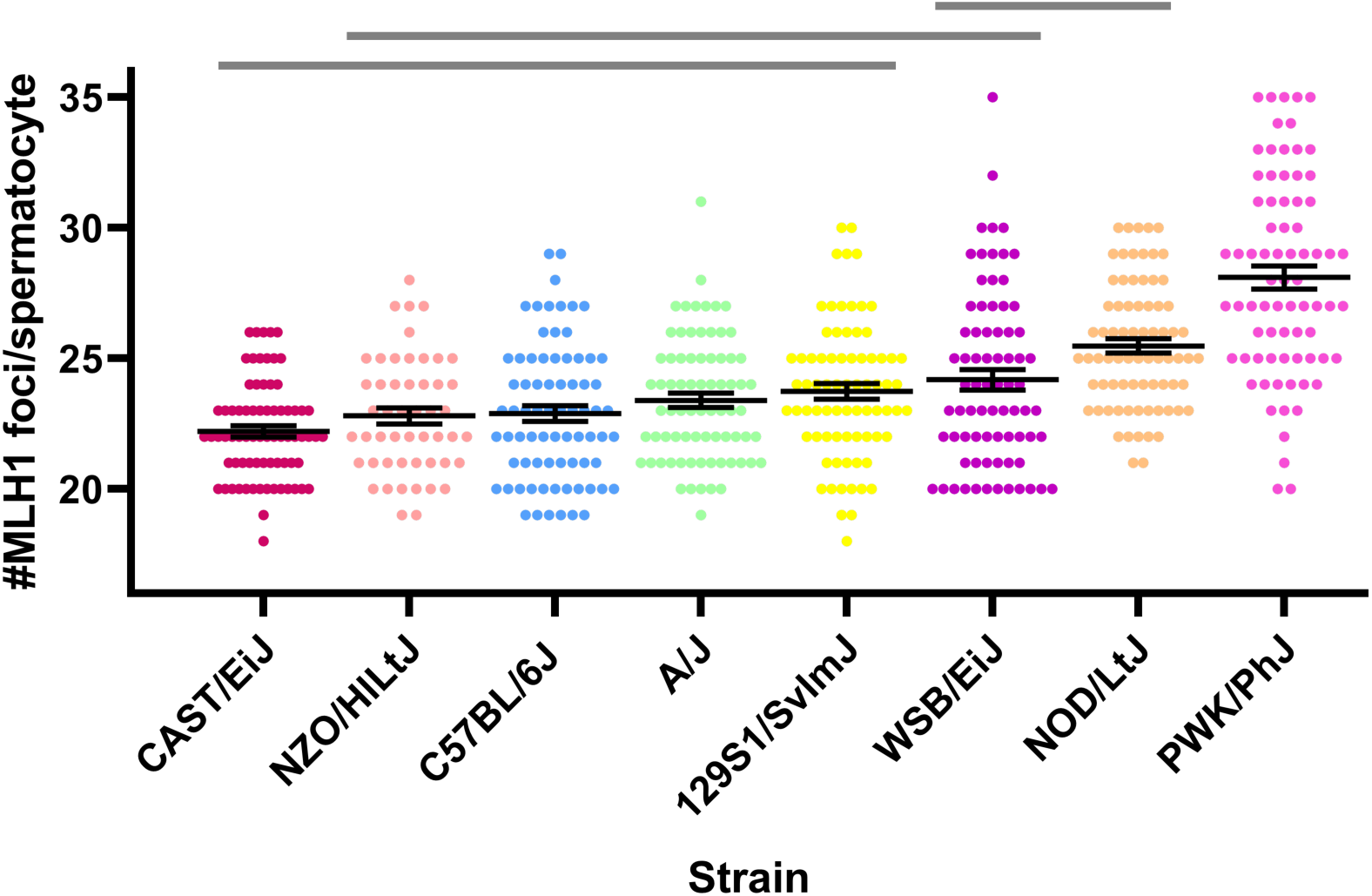
Recombination rate variability among CC founder strains. Autosomal MLH1 foci counts in mid-pachytene spermatocytes are shown for 8 genetically diverse mouse inbred strains, mostly of *M. m. domesticus* origin, except CAST/EiJ and PWK/PhJ, which are *M. m. castaneus* and *M. m. muculus* subspecies, respectively (Yang et al. 2011). Each dot represents the focus count of a single nucleus. Black bars symbolize means ± SEM . Horizontal upper lines represent homogeneous subsets from post hoc comparisons using the Tukey test (α=0.01).

Comparing these with our previous data, we observe that B6 and PWK results are not significantly different than those obtained in our study for the same strains fed with control maintenance diet (*P* = 0.26 and *P* = 0.06, respectively, *t*-test) and are located towards the low and at the high ends of the recombination variability distribution, while MOLF values are intermediate. Our results confirm previous observations of the impact of the genetic background on recombination frequency (Liu et al. 2014) and provide new data about the crossover rate of the CC and DO founder strains.

### Changes in synaptonemal complex length or interference may underlay recombination differences between strains

Crossover distribution and frequency is limited by crossover interference. As a consequence, mouse spermatocytes chromosomes with a short synaptonemal complex (SC) can only undergo one crossover (Lawrie et al. 1995; Petkov et al. 2007; Sym and Roeder 1994; Tease and Hulten 2004). When all chromosomes are considered, cells with longer SCs (measured as µm of immunostained SC) are expected to have more crossovers (Dumont and Payseur 2011a; Froenicke et al. 2002; Kleckner et al. 2003; Lynn et al. 2002). Our data indicate a significant effect of strain on SC length (*P* << 0.0001, ANOVA). When we compare the total length of the SC of the autosomes per cell (in µm, Table 1 and Figure 1B), we observe significantly longer SC in PWK (183.3±19.4) compared to those observed in B6 (162.2±17.1) and MOLF spermatocytes (156.6±16.0; *P* << 0.0001 in both cases, HSD). Hence, the larger SC in PWK may explain the higher crossover frequency observed in this strain respect to those of B6 and MOLF. We found no significant difference in SC length between MOLF and B6 spermatocytes (*P* = 0.20, HSD). This is interesting because these two strains have significant differences in the number of MLH1 foci as reported above. This suggests that factors other than SC length may account for the differences in recombination levels observed between these two strains.

As SC length cannot explain the increase of MLH1 foci in MOLF respect to B6 spermatocytes, we wondered if variation in interference strength could be the cause. As a surrogate for CO interference, we measured the distance between MLH1 foci of bivalents with two or more crossovers. Analysis of variance indicates a significant effect of strain on intercrossover distance (*P* << 0.0001, ANOVA; similar results were obtained with non-parametric tests). Post hoc tests reveal that the average interfocus distance in MOLF spermatocytes is significantly shorter than in B6 and PWK spermatocytes (*P* << 0.0001 both comparisons, HSD; Table 1 and Figure 1C). This is also true when the interfocus distances are expressed as percentage of the length of the SC (de Boer et al. 2009) (*P* << 0.0001 both comparisons, HSD; Table 1 and Figure S2). Our data indicate that not SC length, but interference changes may explain the crossover rate increase observed in MOLF mice respect to that of B6. Therefore, different mechanisms appear to lie beneath the recombination variability between diverse mouse strains.

### Undernourishment may influence recombination levels in a strain-dependent manner

In order to test whether diet could affect recombination, we explored if a 50% restriction to food access in adult males could have an impact on meiotic recombination. Previous studies had shown that a 24-day exposure to dietary changes or environmental factors was sufficient to induce spermatogenesis changes in adult rodents (Assinder et al. 2007; Gely-Pernot et al. 2017). Moreover, a 24-day diet would ensure continuous exposure at least since the spermatogonial stage until pachytene, when recombination is analyzed. Therefore, we chose this time period for our studies about the impact of diets on recombination. We fed adult male mice of each of the three strains with 50% of their regular daily intake of maintenance chow, while controls had access to the same maintenance diet *ad libitum*. At the end of the 24-day treatment, animals were euthanized and testes were processed for crossover analysis. Analysis of the data by a univariate generalized linear model shows that strain (*P* < 0.0001) and diet significantly affect the MLH1 foci frequency (*P*_diet_ = 0.035). We note that 2 of the 3 undernourished PWK animals had to be euthanized before the conclusion of the treatment period due to severe weight loss (20% of body weight). B6 and MOLF were comparatively robust to the effects of undernourishment on body weight. B6 mice often get overweight and can resist well brief periods of food shortage. MOLF is a wild-derived strain of lean mice like PWK, but while the body weight of the latter was reduced 9-20%, MOLF only lost 5-7%.

Neither B6 nor MOLF spermatocytes showed significant changes in MLH1 focus count between *ad libitum* and 50% restricted diets by *post hoc* tests (Table 2), suggesting an efficient control of the recombination levels when adult males of these two strains face undernutrition. In contrast, PWK spermatocytes, already with high crossover frequency, had a small but significant increase (*P* = 0.037) in MLH1 foci number when food intake was restricted to 50% (Table 2). Due to the small magnitude of the effect, we could not determine if it was related to SC length or interference changes. Therefore, our results suggest that, in adult male mice, undernourishment affects recombination levels in particular genetic backgrounds.

**Table 2.** Diet effects on MLH1 foci number per spermatocyte. Comparison of the three types of diets within strains reveals significant differences in MLH foci number in B6 and PWK mice (*P*=0.006 and *P*=0.033, respectively, with ANOVA tests). In the B6 strain, significant differences are observed between the breeding diet and both the maintenance and 50% diet (*P*=0.008 and *P*=0.026, respectively, by Tukey’s tests), but not between these two. In the PWK strain, significant differences are only observed between 50% and maintenance diets (*P*=0.037, Tukey’s test, see text for discussion). Data are the result of the analysis of 75 spermatocytes per treatment and strain group, except B6 maintenance and breeding groups, each with 50 cells. Values are shown as mean±SD. Overall, a significant interaction between strain and diet effects is observed (*P*=0.005).

| Strain | B6 | MOLF | PWK |
| --- | --- | --- | --- |
| Maintenance diet | 23.54±1.97<sup>a</sup> | 24.85±2.57<sup>b</sup> | 29.15±2.87<sup>c</sup> |
| 50% diet | 23.79±1.63<sup>a</sup> | 25.01±2.08 | 30.13±1.95<sup>e</sup> |
| Breeding diet | 24.70±2.21<sup>d</sup> | 24.35±1.62 | 29.95±2.41<sup>ce</sup> |

### Diet composition can affect recombination levels in a strain-dependent manner

We wondered if common laboratory diets could have an effect on recombination and, consequently, may confound the results of recombination studies. Two chows are routinely used in our and many other animal facilities, depending on the purpose: animal maintenance or breeding (see Materials and Methods and Table S1). The breeding diet is aimed to support gestation, lactation and growth, and has 2% more protein and 8% additional energy density than the maintenance chow, devised to promote longevity and normal body weight. In addition, phytoestrogens are present in the breeding diet (150-250 mg isoflavones/kg diet), while avoided in the maintenance one. As previously discussed, these components have been linked to epigenetic or developmental changes in the germline. Hence, we decided to test if crossover frequency could vary in mouse spermatocytes depending on the diet of choice.

Adult mice are routinely fed with maintenance diet. We separated animals of the three strains and provided them with *ad libitum* access to the breeding diet during 24 days, while others were kept with the maintenance diet. After the 24-day period, spermatocytes were obtained for recombination analysis by immunohistochemistry. In order to examine which factors are associated with recombination variability, we used a univariate generalized linear model. Our results indicate that diet significantly affects the MLH1 focus frequency (*P*_diet_ = 0.042). We used post hoc tests to determine which diet comparisons were of particular note statistically. We observed no significant effect of the diets on MLH1 foci frequency in MOLF and PWK mice (Table 2). However, a significant increase was observed in B6 mice fed with breeding diet (24.70±2.21) compared to maintenance chow (23.54±1.97, *P* = 0.008, Table 2). Our results also indicate that strain significantly affects MLH1 frequency (*P* << 0.0001), and there is a significant strain by diet interaction effect (*P* = 0.010) as well. These data therefore indicate that, in addition to genetic differences in recombination frequency, diet composition can affect crossover frequency in adult male mice in a strain-dependent manner.

### Diet effects on recombination are robust to method of analysis

To test whether our findings were robust to the method of statistical analysis, we analyzed these data in aggregate. That is, we used a generalized linear model on data from control, maintenance and calorie restricted diets to test for effects of strain, diet, animal, and any interaction effects. Our results indicate that our findings are robust to statistical approach, with a strong effect of strain (*P* << 0.0001) and a modest but significant effect of diet (*P* = 0.01). We also find an effect of animal (P < 0.001). This model accounts for 62% of phenotypic variance in recombination rate, with the bulk of the variance being accounted for by between strain variation. This is consistent with previous work (*e.g.,* Dumont and Payseur 2011). Importantly, the proportion of the variance due to within-animal sampling is approximately 6%, which indicates in part the consistency of our approach for scoring MLH1 foci.

### Diet effects on recombination levels in C57BL/6 mice are reproducible

We were surprised to find that common chows could affect crossover frequency in B6 mice, while not in the two other strains. In order to find if this was a fortuitous or a consistent observation, we designed a larger experiment (see Materials and Methods). Animals were also subject to maintenance or breeding diet for 24 days and spermatocytes were prepared for analysis immediately after. As shown in Table 3 and Figure 3A, B6 mice kept in maintenance diet had similar MLH1 foci number (23.50±2.17) to those previously observed, and the breeding diet also elicited a significant increase in crossover frequency (24.24±2.31, two-sided Student t-test, *P* = 0.001). This increase could not be explained by significant changes in SC total length (Table 3).

**Figure 3.**
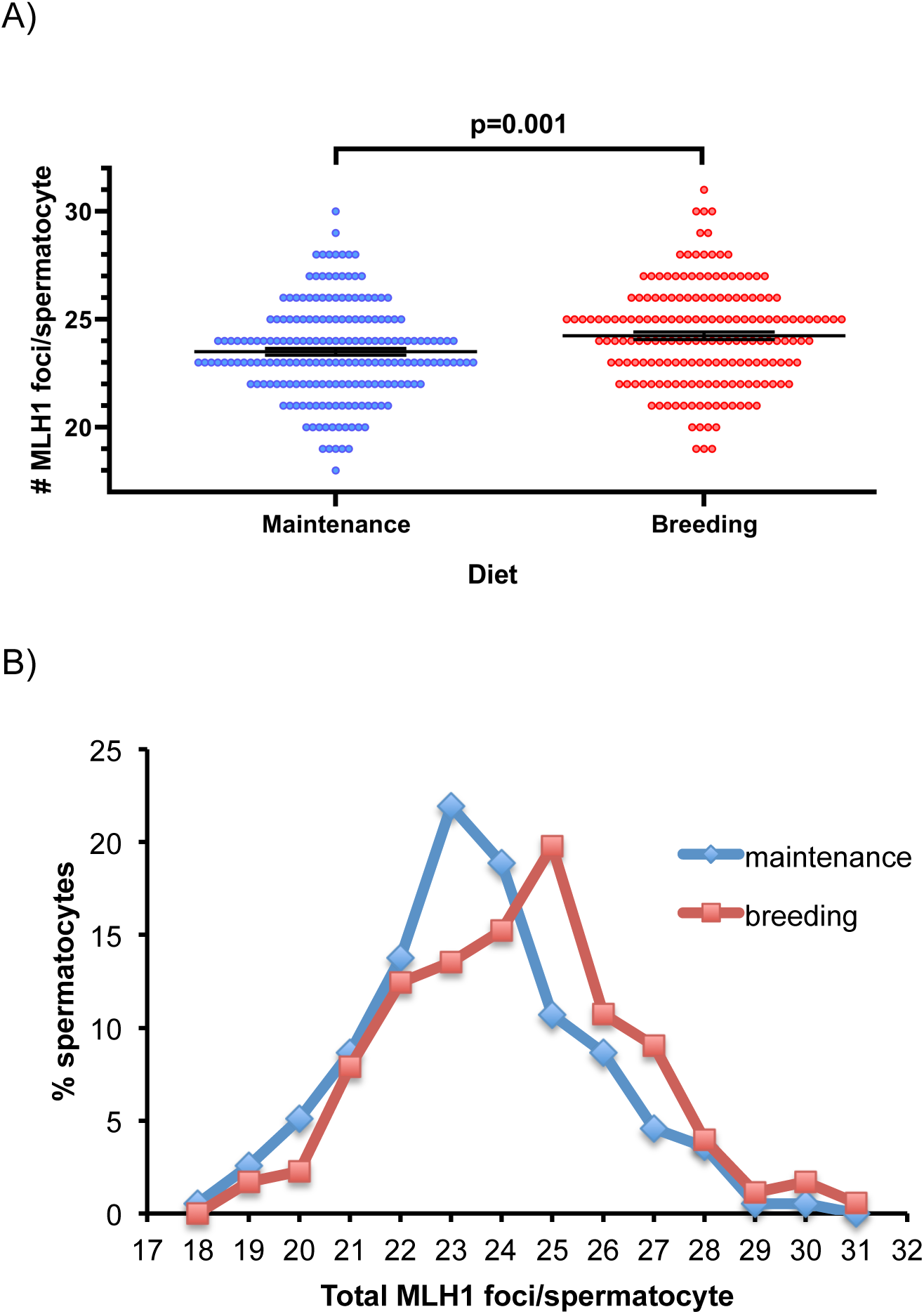
Diet effects on recombination rates. A) Diet effects on recombination rates are strain-dependent. Only C57BL/6 mice show significant differences in recombination rates when fed with maintenance *vs.* breeding diets (see Table 3). Each dot represents the focus count of a single nucleus. Black bars represent means ±SEM. B) Spermatocyte distribution according to the total number of MLH1 foci in C57BL/6 males subject to maintenance and breeding diets.

**Table 3.** Diet effects on crossover frequency in C57BL/6 male mice: analyses per spermatocyte and per bivalent. A) After analyzing an independent group of B6 mice, a significant increase in MLH1 foci count per spermatocyte was again observed in animals fed with breeding diet respect to those kept in maintenance diet (ab, t-student test, *P* = 0.001), while no significant differences in SC length were detected (*P* = 0.20, Mann-Whitney U test). Data are the result of the analysis of 196 spermatocytes in the maintenance group and 177 in the breeding group. B) When bivalents were classified according to the number of crossovers (E0-E3), a significant change was also observed (*P* = 0.004, χ^2^ test). This change was mainly due to an increase of E2 bivalents at the expense of E1 in the breeding group, respect to the maintenance group.

| A) B6 spermatocytes: | Maintenance diet | Breeding diet |
| --- | --- | --- |
| MLH1 foci: | 23.50±2.17<sup>a</sup> | 24.24±2.31<sup>b</sup> |
| SC length (μm): | 169.0±21.2 | 164.6±17.9 |
| B) Number and type of bivalents: |  |  |
| E0: | 19 (0.5%) | 16 (0.5%) |
| E1: | 2808 (75.4%) | 2417 (71.8%) |
| E2: | 895 (24.0%) | 927 (27.5%) |
| E3: | 2 (0.1%) | 5 (0.1%) |

The studies of BPA effects on recombination stemmed from the observation of an increment in chromosome missegregation and aneuploidy caused by this endocrine disruptor (Hunt et al. 2003). In mouse as in other species, proper chromosome segregation requires a minimum of one crossover per chromosome; otherwise, crossover failure often results in aneuploidy (Hassold and Hunt 2001). Hence, we decided to explore if the observed diet effect on recombination had any impact on the frequency of non-recombinant bivalents (E0). We compared the number of bivalents with zero, one, two or three MLH1 foci (E0, E1, E2 and E3, respectively) of each diet group. In both cases, 95% of the spermatocytes had between 20 and 28 MLH1 foci (Figures 3A and 3B) mainly located in E1 and E2 bivalents; those with no crossovers were very rare, as were bivalents with three crossovers, representing 0.5% or less each (Table 3). A significant change was observed between the two diet groups, mainly due to an increase of E2 bivalents at the expense of E1 in animals fed with breeding diet respect to those kept in maintenance diet (χ^2^ = 13.2; *P* = 0.004, Table 3 and Figure 3B). Therefore, we conclude that, compared to the maintenance diet, the breeding chow induces an increase in MLH1 frequency through a shift of E1 to E2 bivalents, without substantial changes in E0 and the associated risk of aneuploidy.

### Diet effects on C57BL/6 spermatogenesis: sperm motility, but not other phenotypes, are affected by diet

We wondered if recombination *per se* was particularly sensitive to dietary changes, or if these were a byproduct of large spermatogenesis disturbances caused by the switch between common diets. Hence, we explored if diet could also affect other aspects of spermatogenesis that resulted in changes in the quality and number of sperm, as observed in several diet and estrogenic exposure studies (Assinder et al. 2007; Horan et al. 2017; Horan et al. 2018; Meena et al. 2017; Nassan et al. 2018; Nätt et al. 2019; Spearow et al. 1999). The breeding diet is more caloric than the maintenance diet (Table S1) and, accordingly, we observed a slight but not significant increase in body and testis weight in animals fed with breeding chow respect to those kept in maintenance diet (Table S2). This small difference disappears when testis weight is corrected for body weight (Table S2). We examined if testes histology was affected by diet, as observed in animals treated with natural or synthetic estrogens (Horan et al. 2017; Spearow et al. 1999). We did not observe noticeable changes in the number and cell composition of seminiferous tubules between the two diet groups (Figure S3). When sperm isolated from the cauda epididymis was analyzed, no differences were detected in sperm count between the two diet groups. Sperm viability was neither significantly affected (Table S2).

Sperm acquires progressive motility in the epididymis, characterized by high velocities and symmetrical, low-amplitude flagellar bends. By computer-assisted sperm analysis (CASA), we assessed several parameters of epididymal sperm motility: percentage of total motility and of progressively motile spermatozoa, VAP, VCL and VSL, LIN, STR, WOB, ALH and BCF (see Methods) (Boyers 1989; Mortimer 1997). None of them showed significant differences between the two diet groups, except the proportion of sperm with progressive motility, which was significantly reduced in the breeding diet (18.8±2.0) respect to the maintenance diet (25.8±2.4, mean±SE; *P* = 0.038) (Table S2). Though not significantly, sperm velocity also appeared to decrease in the breeding diet group (measured as VAP, VCL and VSL). The proportion of motile sperm has been associated with fertilization success (Davis et al. 1991).

In addition, we evaluated whether diet had an effect on sperm DNA integrity by SCSA (Evenson et al. 1980). We analyzed both the DNA fragmentation index (average and total, see Material and Methods), as well as the high DNA stainability; the latter offers a measure of the condensation degree of the sperm chromatin and the percentage of immature sperm cells, because this high stainability is considered to be the result of a lack of full protamination and, thus, an increased histone retention (Evenson et al. 2000). None of these parameters showed significant differences between sperm of mice fed with maintenance *vs.* breeding diets (Table S2). Therefore, these diets had no significant effect on sperm DNA damage or condensation, as measured by SCSA. Only sperm motility was significantly affected by diet of all the phenotypes assayed in sperm and testes.

## DISCUSSION

Sperm quality has declined over the last decades among healthy men (Carlsen et al. 1992; Levine et al. 2017; Sengupta et al. 2017; Splingart et al. 2012). This decline has been associated to chemical exposures and life-style changes, including diets and the increase of diet-related diseases such as obesity (Nassan et al. 2018; Nordkap et al. 2012). Changes in sperm count and motility have a direct impact on fertility. But changes in other underrated sperm features, such as meiotic recombination, are also observed in infertile men (Ferguson et al. 2009; Ren et al. 2016). Alterations of crossover number or distribution increase the risk of chromosome missegregation and aneuploidy (Hassold and Hunt 2001). Hence, fertility requires a tight control of recombination (Cole et al. 2012; Coop and Przeworski 2007).

In spite of this control, several studies have shown that certain environmental factors are able to modify the recombination rate (Belyaev and Borodin 1982; Bomblies et al. 2015; Lloyd et al. 2018; Plough 1917; Singh 2019; Susiarjo et al. 2007; Vrooman et al. 2015). Even if these recombination changes may not be large enough to compromise fertility, they could have important consequences in the transmission and evolution of traits, as well as in genetic mapping studies (Dumont and Payseur 2008; Krzywinska et al. 2016; Pardo-Manuel de Villena et al. 2000; Ritz et al. 2017). To date, only one analysis in flies has reported an effect of nutrition on crossover rate (Mostoufi 2021; Neel 1941), an effect that has also been suggested in yeast (Abdullah and Borts 2001b). Hence, we decided to explore whether diet could not only affect sperm features, but also recombination in mammalian spermatocytes.

### Recombination rate variation among mouse inbred strains: crossover frequency is regulated by different mechanisms

Because both crossover rate and the effect of environmental exposures on recombination depend on the genetic background (Baier et al. 2014; Dumont and Payseur 2011b; Koehler et al. 2002; Vrooman et al. 2015), we selected for our study genetically diverse strains representative of three *Mus musculus* subspecies. We found that crossover frequency, measured by MLH1 immunostaining, was significantly different among them. We reported for the first time very high values in PWK mice, only comparable to those of the PWD/PhJ strain (29.92±2.51 (Dumont and Payseur 2011b); 29.58 (95%CI 28.6630.56) (Balcova et al. 2016)), also of *M. m. musculus* origin (Gregorova and Forejt 2000). B6 values were the lowest and similar to those obtained in previous studies (Baier et al. 2014; Balcova et al. 2016; Vrooman et al. 2014).

For a broader view of recombination variability in mouse, we expanded our analysis to characterize the crossover frequencies of the 8 founder strains of the Collaborative Cross (CC) and Diversity Outbred (DO) stock, because they capture nearly 90% of the known variation present in laboratory mice (Churchill et al. 2004; Roberts et al. 2007). Again, our results showed that PWK spermatocytes have the highest crossover frequency, while CAST/EiJ mice are at the opposite extreme, in agreement with the low recombination levels previously detected in this strain (Baier et al. 2014). Our observations confirm the importance of the genetic background on the levels of recombination (Baier et al. 2014; Dumont 2017; Dumont and Payseur 2011b; Koehler et al. 2002; Liu et al. 2014), and provide new data about the relative crossover rate of the CC and DO founder strains, which are relevant for future mapping and recombination studies in mice.

These results also revealed that the three strains selected for our study represent the low (B6), medium (MOLF) and high (PWK) levels of recombination present in *Mus musculus*. *M. m. molossinus* is considered the result of the hybridization between *M. m. castaneus* and *M. m. musculus*, and while strains of these two subspecies (CAST and PWK) have the lowest and highest recombination rate of all those analyzed in this study, we and others observe an intermediate recombination rate in MOLF (Peterson and Payseur 2021; Silver 1995). Because crossover distribution and frequency depend on crossover interference and chromosome physical length (measured as synaptonemal complex length) (Dumont and Payseur 2011a; Froenicke et al. 2002; Kleckner et al. 2003; Lynn et al. 2002; Petkov et al. 2007; Tease and Hulten 2004), we explored which of these two factors was involved in the recombination differences observed between strains. Moreover, as interference limits the proximity between crossovers, the number of these can increase by either reducing interference or expanding the SC length and we found that both possibilities occurred in the strains under study.

The B6 total autosomal SC length we observe is similar to that previously reported (Vranis et al. 2010). But in PWK, our data suggest that high levels of recombination could be a consequence of the longer SC respect to the other two strains, consistent with the positive correlation between total SC length and recombination rate observed by others in mouse and other animals (Baier et al. 2014; Lynn et al. 2002; Ruiz-Herrera et al. 2017). Interestingly, a recent study has identified several loci that affect SC length and some of them also modulate recombination rate (Wang et al. 2019). A previous study proposed a simple linear relationship between crossover rate and total SC length, so that the ratio between SC length and the number of MLH1 foci would be almost constant in mouse spermatocytes (Lynn et al. 2002). The values observed in our B6 animals coincide with those reported in that study (6.9 µm SC/MLH1 foci; Table 1). However, lower values are observed for MOLF and PWK (6.3 µm), which have higher crossover rates. Wang et al. (2019) and others (Vranis et al. 2010) also found the length of SC per MLH1 focus varies among mouse strains and proposed that it could be a consequence of interference variation, but our PWK data suggest this ratio can change independently of interference fluctuations.

In contrast, differences in total SC length cannot explain the intermediate level of recombination found in MOLF spermatocytes. In this case, the shorter intercrossover distance suggests that, compared to B6, a weaker positive interference in MOLF spermatocytes could be the cause of their higher crossover rate. An inverse correlation between interference strength and recombination rate has also been observed in other mammals (Segura et al. 2013) and a locus that affects both interference and recombination levels has been identified in cattle (Wang et al. 2016). Therefore, our results suggest that diverse and, at least to some extent, independent mechanisms determine the breadth of recombination levels present in mice.

### Recombination rate is both sensitive and resistant to diets: genetic background determines crossover frequency, even under stressful nutritional conditions

Next, we explored if diets can alter recombination levels and whether this effect can be modulated by the genetic background. Based on the findings of the best-studied environmental effects on recombination in mice, those of BPA exposure (Susiarjo et al. 2007; Vrooman et al. 2015), we looked for candidate diets that could also affect the male germline epigenome or compromise sperm function (Manikkam et al. 2013; Rahman et al. 2015; Rahman et al. 2020; Salian et al. 2009; Susiarjo et al. 2013; Xin et al. 2015). Among them, we found that in utero 50% dietary restriction could affect sperm methylation and offspring health (Martínez et al. 2014; Radford et al. 2014). Sperm function and offspring health were also altered by direct undernutrition of adult mouse males (McPherson et al. 2016). When we analyzed if the spermatocytes of males temporarily subjected to undernutrition experienced recombination changes, we found a strain-dependent diet effect.

On the one hand, PWK spermatocytes recombination rate, already high, increased even more when food intake was limited to 50%. Nutritional deficit also causes an increase in crossover frequency in *D. melanogaster* and *S. cerevisiae* (Abdullah and Borts 2001a; Neel 1941). However, we cannot conclude that a reduction in nutrients availability was directly responsible for the observed effect on recombination. Although diverse dietary restrictions have been reported to improve life span (Fontana et al. 2010; McCay et al. 1935), not all mouse strains respond the same; on the contrary, health deterioration and life shortening occurs in some (Liao et al. 2010; Mitchell et al. 2016; Radford et al. 2014). Similarly, we found that while B6 and MOLF animals performed well under reduced food intake, some PWK animals had to be euthanized prematurely due to severe weight loss. Hence, undernutrition may have generated extreme physiological or metabolic disturbances in PWK males capable of altering the control of the recombination levels. Indeed, diverse types of stress have been reported to affect recombination rate in mouse and other organisms (Belyaev and Borodin 1982; Modliszewski and Copenhaver 2017). Though it is possible that meiosis and recombination might have evolved to be able adapt to environmental challenges (Bomblies et al. 2015; Modliszewski and Copenhaver 2017), the adaptive significance of the environment-induced changes in recombination remains unclear. In Arabidopsis, studies on temperature effects on recombination often report U-shaped response curves which suggest that, in optimal conditions, organisms generally have lower recombination rates than under extreme and stressful ones (*e.g.* Lloyd *et al.,* 2018); previous reports of increased recombination rate in stressed mice (Belyaev and Borodin 1982) and our observations are consistent with this prediction. Future studies with different dietary restriction regimes or deprivation of specific nutrients (Ideraabdullah and Zeisel 2018) will be able to discriminate whether food intake reduction or undernourishment-induced stress and health deterioration are responsible for the observed effect on crossing over frequency.

On the other hand, recombination rate in the other two strains (MOLF and B6) was not significantly affected by 50% dietary restriction. Hence, we conclude that, in certain genetic backgrounds, recombination levels are tightly controlled even under stressful conditions such as undernutrition or, as reported in other studies, infections (Dumont et al. 2015). As many organisms, including (unfortunately) many humans, face short, seasonal or long periods of nutrients deprivation, understanding the effects of nutritional changes on recombination, a critically meiotic important process, is important. Given how fundamental diet is to organismal fitness and function, understanding the effect to which diet-induced changes in recombination persist across generations is important as well. Moreover, given that the effects of diet are genotype-specific, more work is needed to comprehend the genetic basis of this interaction.

### Common diets can affect male recombination rate in a strain-dependent manner: recombination in mice is more sensitive to environmental exposures than previously expected

We decided to test whether not just toxicants or stressful exposures, but also small differences between common diets, can alter crossover frequencies in adult male mice. Hence, we temporarily fed adult males of the three selected strains with the two chows routinely used in our facility: one specific for breeding periods, which is richer in proteins and energy density than the regular one, used for mice maintenance, in which phytoestrogen-rich plants are absent in order to avoid estrogenic effects that might interfere with some studies. Although they also differ in other nutrients, their compositions are not markedly dissimilar. Hence, we were surprised to find that males of one of the three strains, B6, showed higher recombination rates when fed with the breeding chow than when kept in the maintenance diet. Again, we observed that the diet effect on recombination was strain-dependent, but now affected to a different strain than undernutrition (PWK), demonstrating the importance of genetic differences in the variable response to diverse diets.

Although a genetic background-dependent response to environmental effects on recombination has been reported in diverse species (Vrooman et al. 2015), we were surprised to find that B6, a strain that is insensitive to the effect BPA and other estrogenic substances on male recombination, was precisely the one responsive to our diet differences (Vrooman et al. 2015). However, those results were also unexpected in view of the high estrogenic sensitivity of B6 testes (Spearow et al. 1999) and previous results reporting a B6 female recombination response to BPA (Susiarjo et al. 2007). Moreover, our results (increased recombination in the breeding diet group, which contains phytoestrogens) were in the opposite direction to the reduced crossover frequency observed in CD-1 mouse males after exposure to synthetic estrogenic substances (Vrooman et al. 2015). Hence, we decided to test whether our observation was a spurious result by providing the same dietary regime to an independent and larger group of B6 adult mice. Our results confirmed that crossover frequency is sensitive to small and apparently healthy diet changes.

These results have important implications, especially for recombination studies, although the chows selected for our study are just a small example of the composition variability found among common rodent diets (Ruhlen et al. 2011). We also wondered if the diet-induced recombination changes could also affect meiotic chromosome segregation and aneuploidy studies, as an increase in achiasmatic chromosomes could compromise meiotic disjunction and elevate the aneuploidy rate, as observed with exposures to BPA and other estrogenic compounds (Hassold and Hunt 2001; Hunt et al. 2003). Our studies showed that the occurrence of achyasmate chromosomes was similarly low in both diet groups. Hence, our results predict that aneuploidy rate and subsequent fertility of B6 mice should not be affected by changes in composition between common chows, although we cannot exclude effects with other diets or genetic backgrounds.

We wondered which mechanisms could explain this diet-induced recombination rate change and if this may be associated to total SC length or interference variation, as we observed for strain-dependent diversity in crossover frequencies. Unfortunately, we were unable to discriminate if interference was affected by diet and could explain our observations. While the diet effect on recombination we detect, though reproducible, is small, intercrossover distances are quite variable; consequently, interference analyses by this method are only possible when relatively large effects on recombination are examined, as those observed by strain effects in our study or by mutations in other reports (Roig et al. 2010). But we could analyze if the observed diet effect on recombination was associated to total SC length variation, as described in recombination changes induced by environmental exposures such as temperature in plants (Lloyd et al. 2018; Modliszewski and Copenhaver 2017; Phillips et al. 2015). However, we did not see significant differences in autosomal SC length between the animals subject to breeding *vs.* maintenance diets. Similarly, Vrooman *et al*. (2015) did not detect SC length changes associated to crossover frequency variation caused by estrogenic substances in adult mice, neither SC length could explain all the temperature effects on recombination in Arabidopsis (Lloyd et al. 2018).

### Recombination can be particularly sensitive to dietary changes

In view of the results, a question emerges: are other aspects of meiosis or spermatogenesis affected? To provide an answer, we examined the testes and sperm of the same mice studied for recombination.

Previous studies had reported changes in spermatogenesis progression, sperm count or motility caused by diets differing in fat, protein or phytoestrogen contents among others (Assinder et al. 2007; Cederroth et al. 2010; Eustache et al. 2009; Matuszewska et al. 2020; Morgan et al. 2020; Tavares et al. 2016). In contrast, and although the breeding chow provides more energy than the maintenance one, the composition differences were insufficient for the breeding diet to cause a significant increase in body or testis weight during the time of exposure. Histological analysis of the testes did not reveal any apparent changes in the morphology or cell content of the seminiferous tubules between the treatment groups, suggesting diets changes did not affect spermatogenesis.

Epididymal sperm count, DNA fragmentation and viability were not significantly affected either, as they were not most of the sperm kinematic parameters analyzed. But interestingly, the percentage of progressively motile spermatozoa decreased after mice transferring to breeding diet, suggesting this diet might be optimal during pregnancy or nursing, but not for male fertility (Davis et al. 1991). Our results are in agreement with those of Nätt *et al*. (2019) and others (Assinder et al. 2007; Nassan et al. 2018; Salas-Huetos et al. 2018), suggesting sperm is capable of rapidly responding to diet, even to small changes.

Indeed, the composition differences between the chows under our study are relatively small compared with those analyzed in studies of high-fat or low-protein diets effects in testes (Crisostomo et al. 2019; Matuszewska et al. 2020; Morgan et al. 2020). However, they are comparable to other diet studies such as the one performed by Assinder *et al*. (2007), who observed changes in testes of adult Wistar rats fed during 24 days with low- or high-phytoestrogen diets (112 µg/g and 465 µg/g, respectively, compared to the 150-250 µg/g of the breeding diet object of our study). This study suggests that particular components of common diets could have an impact on spermatogenesis and sperm quality.

Our findings raise the question as to whether the two observed diet-induced effects (on recombination levels in pachytene spermatocytes and on epidydimal sperm motility) are related or not and caused by the same of by different dietary components. BPA studies have revealed effects not only on recombination, but also on sperm motility and at multiple stages of spermatogenesis (Rahman et al. 2015; Tiwari and Vanage 2013; Vrooman et al. 2015). Though the relation between the different effects is unclear, our results show that the switch between common chows does not cause major disturbances in spermatogenesis that could also perturb recombination. On the contrary, recombination can be particularly sensitive to dietary changes.

### Open questions and implications for recombination studies

What is the mechanism that links diet with recombination? It is tempting to speculate that epigenetic changes, such as histone modifications and DNA methylation, could be involved in the diet effect on recombination rate, because crossover frequency and distribution depend on the chromatin architecture and epigenetic marks of the chromosomes (Buard et al. 2009; de la Casa-Esperon and Sapienza 2003; Kleckner et al. 2003; Zelkowski et al. 2019 (Termolino et al. 2016). Germline epigenetic modifications have been found to be susceptible to dietary changes (*e.g.,* in energy, protein or phytoestrogen content) and to environmental exposures capable of affecting recombination (*e.g.,* BPA, atrazine) (Gely-Pernot et al. 2017; Manikkam et al. 2013; Modliszewski and Copenhaver 2017; Xin et al. 2015). Indeed, it has been proposed that recombination rate could vary as a consequence of the germline epigenetic response to environmental exposures (Modliszewski and Copenhaver 2017). Epigenetic modifications may also be the underlying cause of the sperm motility differences observed between the two diet groups, as other diets have been reported to elicit both sperm epigenome and motility changes (Nätt et al. 2019; Siddeek et al. 2018).

Although the SCSA results did not suggest large sperm chromatin alterations due to diets, this is not surprising in view of the moderate differences between the diets under study and the reproductive success of the animals fed with them in facilities throughout the world. Moreover, diet-induced epigenetic changes in the male germline have been shown to be heterogeneous among studies (Donkin and Barres 2018; Sharma and Rando 2017; Siddeek et al. 2018); for instance, the nature of the epigenetic marks that result in transgenerational inheritance has been questioned, as they are either of small magnitude, variable sort or even undetectable in some generations (Sharma and Rando 2017; Shea et al. 2015; Xue et al. 2016). Hence, although the sperm epigenome has been proposed as a marker for environmental exposures, the analyses often turn out to be very complicated (Siddeek et al. 2018). In contrast, our results show that crossover rate is sensitive not only to disrupting toxicants (Horan et al. 2018), but also to small changes in diet and could potentially be used as an indicator of environmentally-induced perturbations in the germline.

Our results also show that this recombination sensitivity depends on the genetic background, which is also true for many other responses to diverse exposures, including diets (Latchney et al. 2018; Spearow et al. 1999; Thigpen et al. 2007; Vrooman et al. 2015). For this reason, studies about the effects of environmental factors must explore their impact in genetically diverse strains, such as the founders of the CC and DO mice. Disparate results can also result from variability in the doses, timing and duration of exposure, among others. For instance, the effects of BPA on recombination are developmental stage, sex and strain dependent (Susiarjo et al. 2007; Vrooman et al. 2015). We now add a further factor to control in recombination studies: diet.

Future studies will determine which component of the chows (or combination of them) is responsible for the observed changes in recombination, as well as the effective doses. Phytoestrogens are attractive candidates, as they can elicit germline epigenetic as well as sperm motility changes, have estrogenic properties like BPA and can even modulate the effects of this compound in mice (Atanassova et al. 2000; Dolinoy et al. 2007; Muhlhauser et al. 2009; Patisaul 2017). But changes in energy content or even minor components of the diets have also shown to affect the germline and constitute interesting candidates (Crisostomo et al. 2019; Nassan et al. 2018; Ruhlen et al. 2011; Salas-Huetos et al. 2018; Siddeek et al. 2018).

Finally, while BPA and other estrogenic compounds affect recombination when provided to female embryos or neonatal males (Gely-Pernot et al. 2017; Susiarjo et al. 2007; Vrooman et al. 2015), our results demonstrate that recombination in adult male mice is sensitive to diet influences. It will be interesting to explore whether earlier developmental stages, particularly those in which the germline epigenetic reprogramming takes place and are particularly vulnerable to exposures such as endocrine disruptors (Ly et al. 2015; McCarrey 2014), as well as female recombination, are also susceptible to diet effects.

In conclusion, our study in mice shows that male recombination rate is sensitive to dietary changes, and this sensitivity depends on the genetic background. This is the first report of a diet effect on genome-wide levels of recombination. Our results send a cautionary note for recombination studies, as diet constitutes a new factor that should be taken into account.

## Data availability

The data underlying this article will be shared on reasonable request to the corresponding author.

## Acknowledgments

We would like to thank Dolores García Olmo and Isabel Blanco Gutierrez for donating animals and materials. We acknowledge Julia Maria Samos Juarez for advice with the diets experimental design, as well as the Albacete UCLM Animal Experimentation Center staff for mouse monitoring and feeding according to the experimental procedure; this was possible thanks to the UCLM Vicerrectorado de Investigacion and J. Julian Garde. We are grateful to Jose Ramon Marin Tebar for capturing the microscope images and to Joaquim Soriano Felipe for support with SC automated analysis. We express our gratitude to the Biobank of Albacete for processing the testicular tissue samples. We also would like to thank Matthieu Falque and Olivier C. Martin for discussions about chromosome interference. We gratefully acknowledge Harry Sedgwick, Beth Dumont, and David Threadgill for their assistance with the analysis of the CC mice. E. de la Casa-Esperon received financial support through the program “Plan Propio de Investigacion” of the University of Castilla-La Mancha (2018/11744), co-funded by the European Regional Development Fund (FEDER, UE).

The authors declare that there is no conflict of interest.

