## Supplementary material for "Diet effects on mouse meiotic recombination: a warning for recombination studies": Table S1

**Table S1: Summary of diets composition**

| Chow | Maintenance: Teklad 2014 | Breeding: Harlan Teklad 2018 | LabDiet 5001 |
|---|---|---|---|
| Isoflavones | 0-20 mg/kg | 150-250 mg/kg | high* |
| Protein | 14.3% | 18.6% | 23.9% |
| Fat extract | 4.0% | 6.2% | 5.0% |
| Carbohydrate (available) | 48.0% | 44.2% | n/a |
| Crude fiber | 4.1% | 3.5% | 5.1% |
| Neutral detergent fiber | 18.0% | 14.7% | 15.6% |
| Ash | 4.7% | 5.3% | 7.0 |
| Energy density | 2.9 kcal/g | 3.1 kcal/g | 3.4 kcal/g |
| Calories from protein | 20% | 24% | 28.5% |
| Calories from fat | 13% | 18% | 13.5% |
| Calories from carbohydrate | 67% | 58% | 58.0% |

2014: Teklad Global 14% Protein Rodent Maintenance Diet (Harlan Laboratories), designed to promote longevity and normal body weight in rodents. It does not contain alfalfa or soybean meal in order to minimize the occurrence of natural phytoestrogens. Capsumlab Maintenance Complete Chow (Capsumlab, Spain) has the same composition.

2018: Teklad Global 18% Protein Rodent Diet (Harlan Laboratories), formulated to support gestation, lactation and growth of rodents. It contains soy, but not alfalfa in order to limit the amount of phytoestrogens.

Laboratory Rodent Diet 5001 (LabDiet 5001 or Purina 5001): a classical chow designed for life-cycle nutrition (maintenance), but not for maximizing production in mouse breeding colonies. It contains soy and alfalfa.

Mineral and vitamin content, as well as other detailed information about the composition are also available from the manufacturers websites.

*Thigpen et al. (2013) reported that Purina 5001 contains 484±113 mg/kg daidzein+genistein (810±10 mg/kg according to Brown and Setchell (2001), mostly daidzin and genistein (72%), plus smaller amounts of other conjugates and aglycones). The isoflavone contents reported by Harlan Laboratories refer to daidzein plus genistein aglycone equivalents.

*J. E. Thigpen et al. (2013) The estrogenic content of rodent diets, bedding, cages, and water bottles and its effect on bisphenol A studies. Journal of the American Association for Laboratory Animal Science 52, 130-141.*

*N. M. Brown and K. D. R. Setchell (2001) Animal models impacted by phytoestrogens in commercial chow: implications for pathways influenced by hormones. Laboratory Investigation 51, 735-747*
