## Supplementary material for "Diet effects on mouse meiotic recombination: a warning for recombination studies": Table S2

**Table S2: Diet effects on body and testis weight, sperm kinematic parameters and SCSA measurements.**

| C57BL/6 mice diet: | Maintenance | Breeding | *P*-value |
|---|---|---|---|
| Body weight (g) | 31.5±1.4 | 32.3±1.1 | 0.410 |
| Testis weight (g) | 0.105±0.003 | 0.108±0.002 | 0.410 |
| Testis/body weight (%) | 0.338±0.017 | 0.334±0.006 | 0.799 |
| Sperm count/ml | $2.27 \cdot 10^7 \pm 0.55 \cdot 10^7$ | $2.27 \cdot 10^7 \pm 0.31 \cdot 10^7$ | 0.566 |
| Sperm viability (%) | 59.2±4.1 | 57.3±1.9 | 0.655 |
| tDFI (%) | 4.55±1.07 | 5.71±1.24 | 0.760 |
| HDS (%) | 13.3±5.3 | 12.1±5.6 | 0.759 |
| Progressive motility (%) | 25.8±2.4 | 18.8±2.0 | 0.038* |
| Total motility (%) | 60.5±3.6 | 57.8±3.1 | 0.580 |
| VCL (µm/s) | 88.8±6.3 | 71.3±5.4 | 0.061 |
| VSL (µm/s) | 47.3±4.3 | 36.3±3.6 | 0.064 |
| VAP (µm/s) | 61.5±5.0 | 48.7±4.2 | 0.062 |
| LIN (%) | 51.0±2.3 | 47.5±2.0 | 0.266 |
| STR (%) | 72.2±1.9 | 68.5±1.6 | 0.140 |
| WOB (%) | 67.8±1.7 | 66.5±1.5 | 0.563 |
| ALH (µm) | 3.36±0.20 | 2.84±0.17 | 0.055 |
| BCF (µm) | 6.35±0.28 | 5.76±0.23 | 0.120 |

Testis/body weight: testis weight fraction of body weight. tDFI, total DNA fragmentation index; HDS, high DNA stainability; VCL, curvilinear velocity; VSL, straight line velocity; VAP, average path velocity; LIN, linearity; STR, straightness; WOB, wobble; ALH, lateral head displacement; BCF, beat cell frequency. Body, testis, sperm and SCSA data were analyzed by two-sided Student t-test; values are shown as mean ± SEM. Kinematic parameters were analyzed by factorial ANOVA and values are shown as LS mean ± SE. * $P < 0.05$
