## Supplementary material for "Diet effects on mouse meiotic recombination: a warning for recombination studies": Figure S1

A)

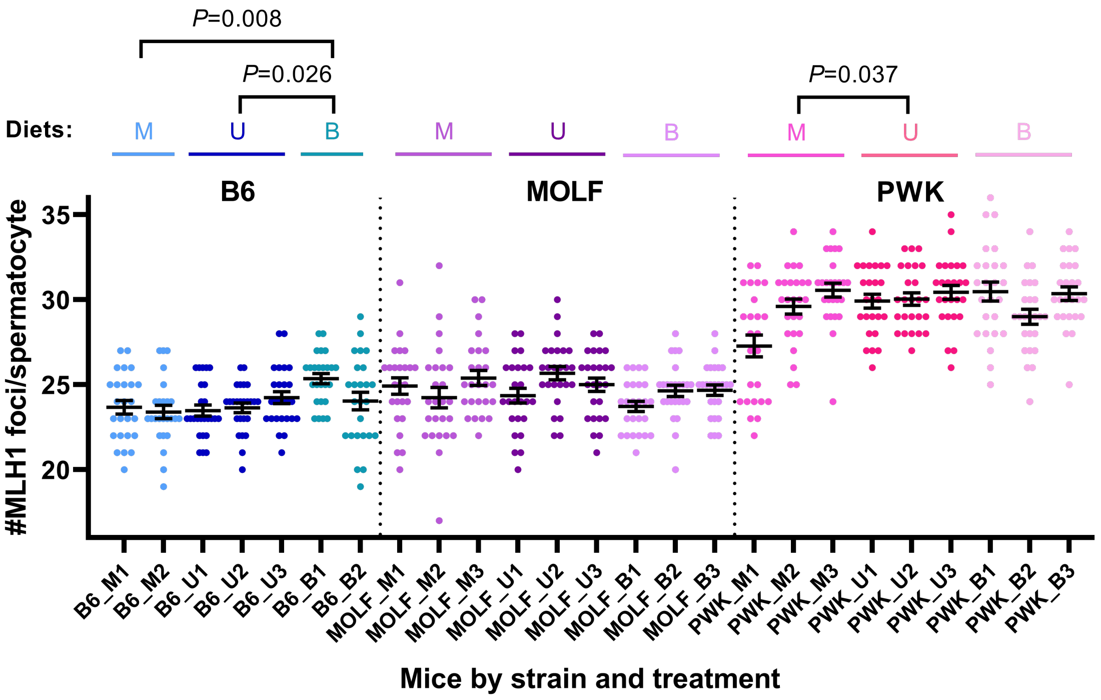


B)

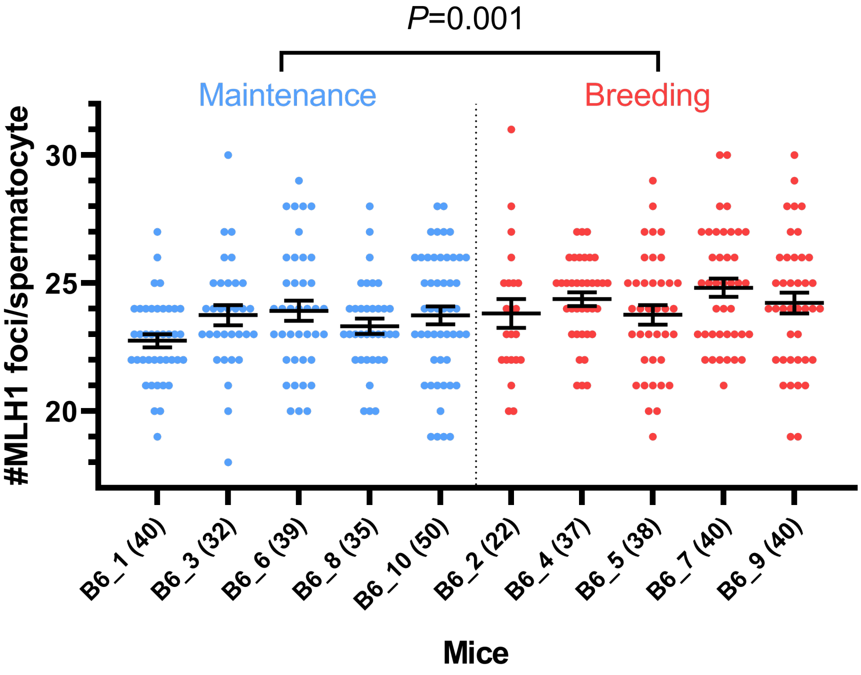


**Figure S1** Strain and diet effects on recombination rate. A) Crossover rates of each animal are grouped by strain (B6, MOLF or PWK) and diet (M=maintenance, U= undernourishment, B= breeding). Numbers identify animals within the same diet and strain group. Data are the result of 25

spermatocytes analyzed per mouse. B) Diet effects on C57BL/6 mice. 5 mice were fed with maintenance chow and 5 with breeding diet. The number of spermatocytes analyzed in each one is indicated in parenthesis. Black bars represent means $\pm$ SEM. *P*-values were calculated for data pooled per strain and diet as described in Tables 2 and 3 and in the main text. As noted in the main text, the effects of strain and diet are robust to the method of statistical analysis.
