## Supplementary material for "Diet effects on mouse meiotic recombination: a warning for recombination studies": Figure S2

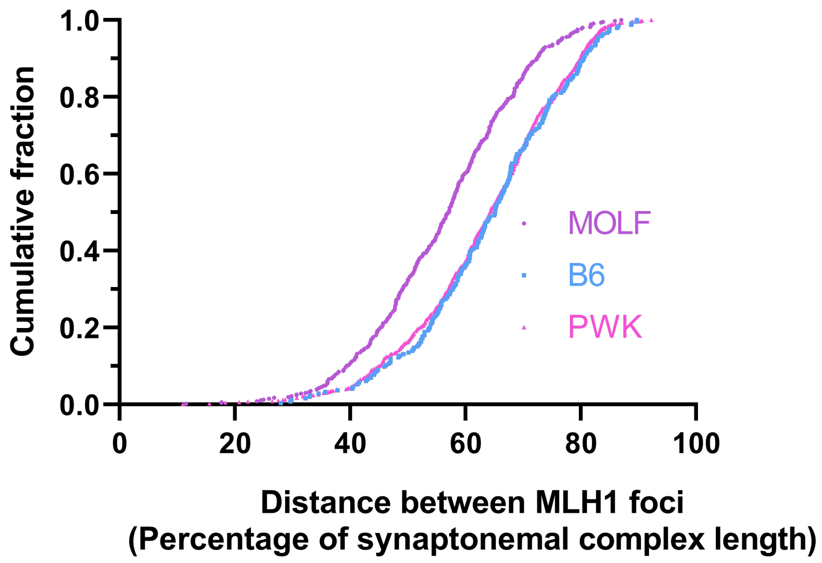


**Figure S2** Strain effect on interference. Cumulative fraction of the intercrossover distances as percentage of synaptonemal complex length, which are significantly shorter in MOLF males respect to the other two strains (Table 1)
