## Supplementary material for "Diet effects on mouse meiotic recombination: a warning for recombination studies": Figure S3

A)

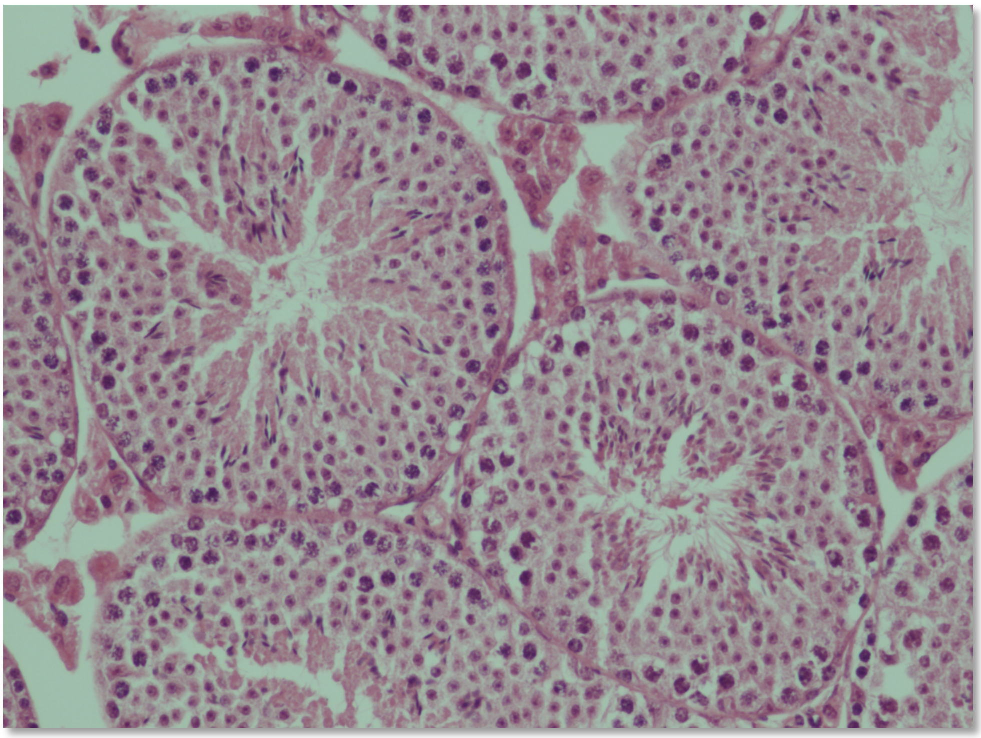

B)

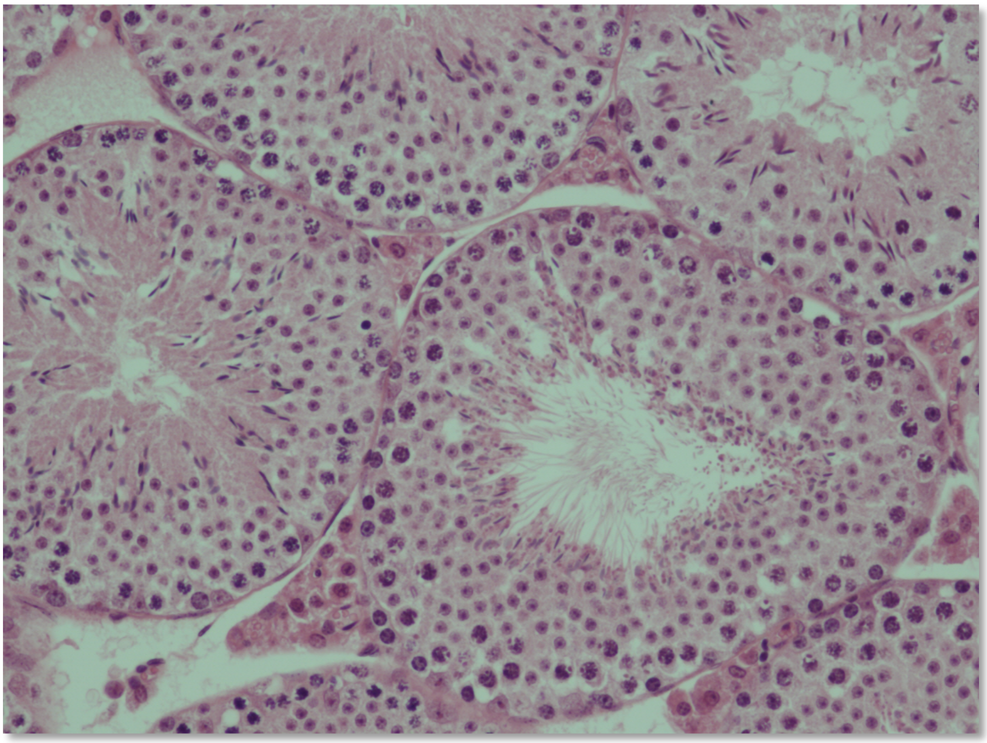

**Figure S3** Histological analysis does not reveal any diet effect on seminiferous tubules cell-type composition. A) Example of testis section of a mouse fed with maintenance diet and B) of breeding diet-treated male, both stained with hematoxylin and eosin.
